# Biogeography & environmental conditions shape bacteriophage-bacteria networks across the human microbiome

**DOI:** 10.1101/144642

**Authors:** Geoffrey D Hannigan, Melissa B Duhaime, Danai Koutra, Patrick D Schloss

**Affiliations:** Department of Microbiology & Immunology, University of Michigan, Ann Arbor, Michigan, 48109, USA; Department of Ecology & Evolutionary Biology, University of Michigan, Ann Arbor, Michigan, 48109, USA; Department of Computer Science, University of Michigan, Ann Arbor, Michigan, 48109, USA

## Abstract

Viruses and bacteria are critical components of the human microbiome and play important roles in health and disease. Most previous work has relied on studying bacteria and viruses independently, thereby reducing them to two separate communities. Such approaches are unable to capture how these microbial communities interact, such as through processes that maintain community robustness or allow phage-host populations to co-evolve. We implemented a network-based analytical approach to describe phage-bacteria network diversity throughout the human body. We built these community networks using a machine learning algorithm to predict which phages could infect which bacteria in a given microbiome. Our algorithm was applied to paired viral and bacterial metagenomic sequence sets from three previously published human cohorts. We organized the predicted interactions into networks that allowed us to evaluate phage-bacteria connectedness across the human body. We observed evidence that gut and skin network structures were person-specific and not conserved among cohabitating family members. High-fat diets appeared to be associated with less connected networks. Network structure differed between skin sites, with those exposed to the external environment being less connected and likely more susceptible to network degradation by microbial extinction events. This study quantified and contrasted the diversity of virome-microbiome networks across the human body and illustrated how environmental factors may influence phage-bacteria interactive dynamics. This work provides a baseline for future studies to better understand system perturbations, such as disease states, through ecological networks.

**Author Summary:** The human microbiome, the collection of microbial communities that colonize the human body, is a crucial component to health and disease. Two major components of the human microbiome are the bacterial and viral communities. These communities have primarily been studied separately using metrics of community composition and diversity. These approaches have failed to capture the complex dynamics of interacting bacteria and phage communities, which frequently share genetic information and work together to maintain ecosystem homestatsis (e.g. kill-the-winner dynamics). Removal of bacteria or phage can disrupt or even collapse those ecosystems. Relationship-based network approaches allow us to capture this interaction information. Using this network-based approach with three independent human cohorts, we were able to present an initial understanding of how phage-bacteria networks differ throughout the human body, so as to provide a baseline for future studies of how and why microbiome networks differ in disease states.

## Introduction

Viruses and bacteria are critical components of the human microbiome and play important roles in health and disease. Bacterial communities have been associated with disease states, including a range of skin conditions [1], acute and chronic wound healing conditions [2,3], and gastrointestinal diseases, such as inflammatory bowel disease [4,5], *Clostridium difficile* infections [6] and colorectal cancer [7,8]. Altered human viromes (virus communities consisting primarily of bacteriophages) also have been associated with diseases and perturbations, including inflammatory bowel disease [5,9], periodontal disease [10], spread of antibiotic resistance [11], and others [12–17]. Viruses act in concert with their microbial hosts as a single ecological community [18]. Viruses influence their living microbial host communities through processes including lysis, host gene expression modulation [19], influence on evolutionary processes such as horizontal gene transfer [22] or antagonistic co-evolution [26], and alteration of ecosystem processes and elemental stoichiometry [27].

Previous human microbiome work has focused on bacterial and viral communities, but have reduced them to two separate communities by studying them independently [5,9,10,12–17]. This approach fails to capture the complex dynamics of interacting bacteria and phage communities, which frequently share genetic information and work together to maintain ecosystem structure (e.g. kill-the-winner dynamics that prevent domination by a single bacterium). Removal of bacteria or phages can disrupt or even collapse those ecosystems [18,28–37]. Integrating these datasets as relationship-based networks allow us to capture this complex interaction information. Studying such bacteria-phage interactions through community-wide networks built from inferred relationships begins to provide us with insights into the drivers of human microbiome diversity across body sites and enable the study of human microbiome network dynamics overall.

In this study, we characterized human-associated bacterial and phage communities by their inferred relationships using three published paired virus and bacteria-dominated whole community metagenomic datasets [13,14,38,39]. We leveraged machine learning and graph theory techniques to establish and explore the human bacteria-phage network diversity therein. This approach built upon previous large-scale phage-bacteria network analyses by inferring interactions from metagenomic datasets, rather than culture-dependent data [33], which is limited in the scale of possible experiments and analyses. We implemented an adapted metagenomic interaction inference model that made some improvements upon previous phage-host interaction prediction models. Previous approaches have utilized a variety of techniques, such as linear models that were used to predict bacteria-phage co-occurrence using taxonomic assignments [40], and nucleotide similarity models that were applied to both whole virus genomes [41] and clusters of whole and partial virus genomes [42]. Our approach uniquely included protein interaction data and was validated based on experimentally determined positive and negative interactions (i.e. who does and does not infect whom). We built on previous modeling work as a means to our ends, and focused on the biological insights we could gain instead of building a superior model and presenting our work as a toolkit. We therefore did not focus on extensive benchmarking against other existing models [41,43–45]. Through this approach we were able to provide an initial basic understanding of the network dynamics associated with phage and bacterial communities on and in the human body. By building and utilizing a microbiome network, we found that different people, body sites, and anatomical locations not only support distinct microbiome membership and diversity [13,14,38,39,46–48], but also support ecological communities with distinct communication structures and robustness to network degradation by extinction events. Through an improved understanding of network structures across the human body, we aim to empower future studies to investigate how these communities dynamics are influenced by disease states and the overall impact they may have on human health.

## Results

### Cohort Curation and Sample Processing

We studied the differences in virus-bacteria interaction networks across healthy human bodies by leveraging previously published shotgun sequence datasets of purified viral metagenomes (viromes) paired with bacteria-dominated whole community metagenomes. Our study contained three datasets that explored the impact of diet on the healthy human gut virome [14], the impact of anatomical location on the healthy human skin virome [13], and the viromes of monozygotic twins and their mothers [38,39]. We selected these datasets because their virome samples were subjected to virus-like particle (VLP) purification, which removed contaminating DNA from human cells, bacteria, etc. To this end, the publishing authors employed combinations of filtration, chloroform/DNase treatment, and cesium chloride gradients to eliminate organismal DNA (e.g. bacteria, human, fungi, etc) and thereby allow for direct assessment of both the extracellular and fully-assembled intracellular virome (**Supplemental Figure S1 A-B**) [14,39]. Each research group reported quality control measures to ensure the purity of the virome sequence datasets, using both computational and molecular techniques (e.g. 16S rRNA gene qPCR) (**Table S1**). These reports confirmed that the virome libraries consisted of highly purified virus genomic DNA.

The bacterial and viral sequences from these studies were quality filtered and assembled into contigs (i.e. genomic fragments). We further grouped the related bacterial and phage contigs into operationally defined units based on their k-mer frequencies and co-abundance patterns, similar to previous reports (**Supplemental Figure S2-S3**) [42]. This was done both for dimensionality reduction and to prevent inflation of node counts by using contigs which are expected to represent multiple fragments from the same genomes. This was also done to create genome analogs that we could use in our classification model which was built using genome sequences. We referred to these operationally defined groups of related contigs as operational genomic units (OGUs). Each OGU represented a genomically similar sub-population of either bacteria or phages. Contig lengths within clusters ranged between 10^3^ and 10^5.5^ bp (**Supplemental Figure S2-S3**).

The original publications reported that the whole metagenomic shotgun sequence samples, which primarily consisted of bacteria, had an average viral relative abundance of 0.4% (**Table S1**) [13,14,38,39]. We confirmed these reports by finding that only 2% (6 / 280 OGUs) of bacterial OGUs had significantly strong nucleotide similarity to phage reference genomes (e-value < 10^−25^) [13,14,38,39]. Additionally, no OGUs were confidently identified as lytic or temperate phage OGUs in the bacterial dataset using the Virsorter algorithm [50]. We also supplemented the previous virome fraction quality control measures (**Table S1**) to find that, in light of the rigorous purification and quality control during sample collection and preparation, 77% (228 / 298 operational genomic units) still had some nucleotide similarity to a given bacterial reference genome (e-value < 10^−25^). It is important to note that interpreting such alignment is complicated by the fact that most reference bacterial genomes also contain prophages (i.e. phages integrated into bacterial genomes), meaning we do not know to what extent the alignments were the result of bacterial contaminants in the virome fraction and what were true integrated prophages. As most phages in these communities have been shown to be temperate (i.e. they integrate into bacterial genomes) using methods that included nucleotide alignments of phages to bacterial reference genomes, we expected that a large fraction of those phages were temperate and therefore shared elements with bacterial reference genomes – a trend previously reported [14]. To ensure the purity of our sample sets, we supplemented the quality control measures by filtering out all OGUs that could be potential bacterial contaminants, as described previously [42]. This resulted in the removal of 143 OGUs that exhibited nucleotide similarity to bacterial genomes but no identifiable known phage elements. We were also able to identify two OGUs as representing **complete**, high confidence phages using the stringent Virsorter phage identification algorithm (class 1 confidence group) [50].

### Implementing Phage-Bacteria Interaction Prediction to Build a Community Network

We predicted which phage OGUs infected which bacterial OGUs using a random forest model trained on experimentally validated infectious relationships from six previous publications [41,51–55]. Only bacteria and bacteriophages were used in the model. The training set contained 43 diverse bacterial species and 30 diverse phage strains, including both broad and specific ranges of infection (**Supplemental Figure S4 A-B, Table S2**). While it is true that there are more known phages that infect bacteria, we were limited by the information confirming which phages do not infect certain bacteria and attempted to keep the numbers of positive and negative interactions similar. Phages with linear and circular genomes, as well as ssDNA and dsDNA genomes, were included in the analysis. Because we used DNA sequencing studies, RNA phages were not considered (**Supplemental Figure S4 C-D**). This training set included both positive relationships (i.e. a phage infects a bacterium) and negative relationships (i.e. a phage does not infect a bacterium). This allowed us to validate the false positive and false negative rates associated with our candidate models, thereby building upon previous work that only considered positive relationships [41]. It is also worth noting that while a positive interaction is strong evidence that the interaction exists, we must also be conscious that negative interactions are only weak evidence for a lack of interaction because the finding could be the result of our inability to reproduce conditions in which those interactions occur. Altogether we decided to maintain a balanced dataset at the cost of under-sampling the available positive interaction information because the use of such a severely unbalanced dataset often results in over-fit and uninformative model training. However, as an additional validation measure, we used the extensive additional positive interactions as a secondary dataset to confirm that we could identify infectious interactions from a more diverse bacterial and phage dataset. Using this approach, we confirmed that 382 additional phage reference genomes, representing a diverse range of phages, were matched to at least one reference bacterial host genome of the species that they were expected to infect (**Supplemental Figure S5**). Because the model was built on full genomes and used on OGUs, we also assessed whether our model was resilient to incomplete reference genomes. We found that the use of our model on random contigs representing as little as 50% length of the original reference phage and bacterial genomes resulted in minimal reduction in the ability of the model to identify relationships (**Supplemental Figure S6**).

Four phage and bacterial genomic features were used in our random forest model to predict infectious relationships between bacteria and phages: 1) genome nucleotide similarities, 2) gene amino acid sequence similarities, 3) bacterial Clustered Regularly Interspaced Short Palindromic Repeat (CRISPR) spacer sequences that target phages, and 4) similarity of protein families associated with experimentally identified protein-protein interactions [56]. These features were calculated using the training set described above. While the nucleotide and amino acid similarity metrics were expected to identify prophage signatures, the protein family interaction and CRISPR signatures were expected to aid in identifying lytic phages in addition to temperate phages. We chose to utilize these metrics that directly compare nucleotide sequences between sample phages and bacteria, instead of relying on alignment to reference genomes or known marker genes, because we were extrapolating our model to highly diverse communities which we expect to diverge significantly from the available reference genomes. The resulting random forest model was assessed using nested cross validation, and the median area under its receiver operating characteristic (ROC) curve was 0.788, the median model sensitivity was 0.905, and median specificity was 0.538 (**Figure 1 A**). This balance of confident true positives at the cost of fewer true negatives was ideal for this type of dataset which consisted of primarily positive connections (**Supplemental Figure S7**). Nested cross validation of the model demonstrated that the sensitivity and specificity of the model could vary but the overall model performance (AUC) remained more consistent (**Supplemental Figure S8**). This suggested that our model would perform with a similar overall accuracy despite changes in sensitivity/specificity trade-offs. The most important predictor in the model was amino acid similarity between genes, followed by nucleotide similarity of whole genomes (**Figure 1 B**). Protein family interactions were moderately important to the model, and CRISPRs were largely uninformative, due to the minimal amount of identifiable CRISPRs in the dataset and their redundancy with the nucleotide similarity methods (**Figure 1 B**). Approximately one third of the training set relationships yielded no score and therefore were unable to be assigned an interaction prediction (**Figure 1 C**).

**Figure 1:**
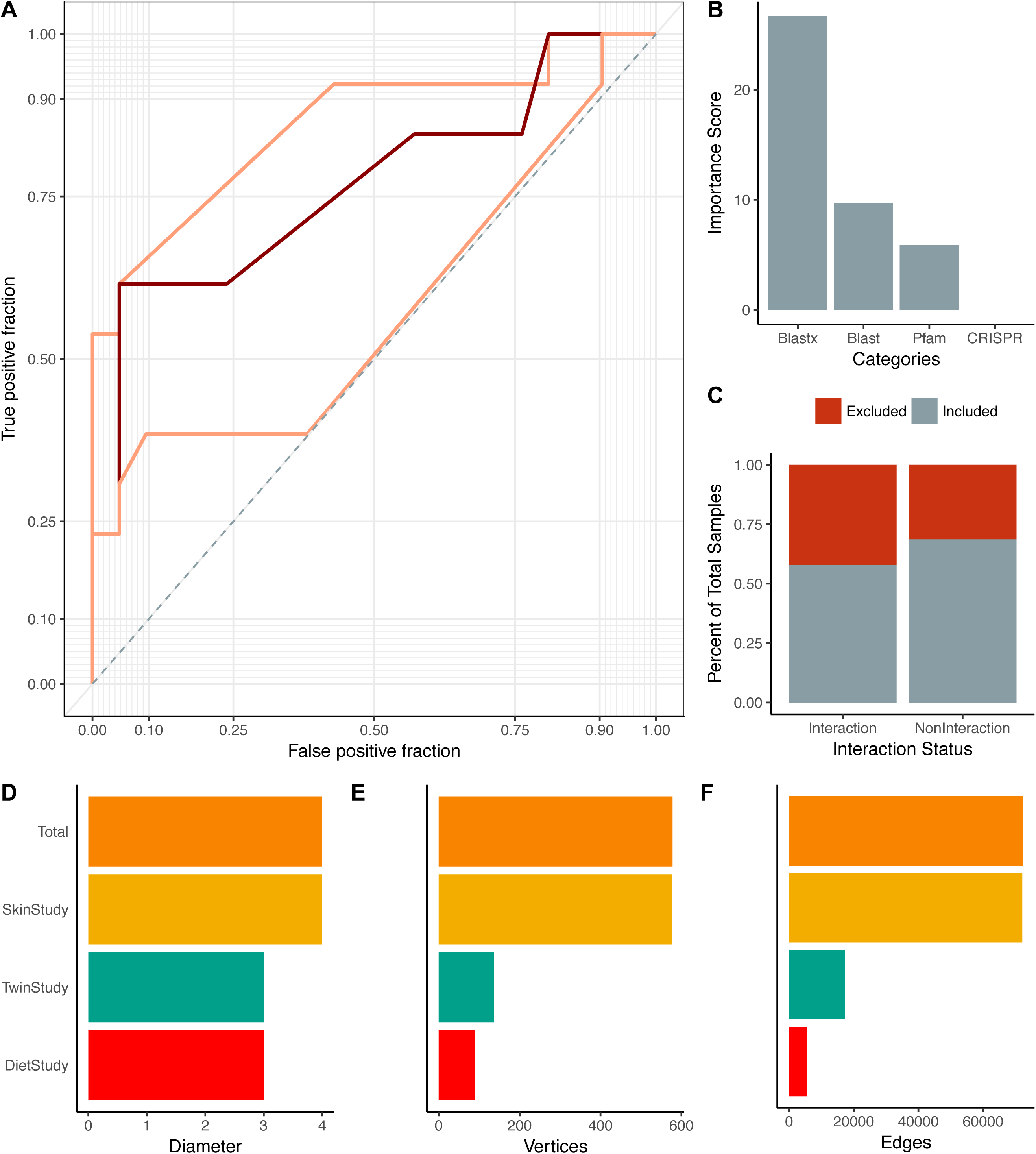
Summary of Multi-Study Network Model. (A) Median ROC curve (dark red) used to create the microbiome-virome infection prediction model, based on nested cross validation over 25 random iterations. The maximum and minimum performance are shown in light red. (B) Importance scores associated with the metrics used in the random forest model to predict relationships between bacteria and phages. The importance score is defined as the mean decrease in accuracy of the model when a feature (e.g. Pfam) is excluded. Features include the local gene alignments between bacteria and phage genes (denoted `blastx`; the blastx algorithm in Diamond aligner), local genome nucleotide alignments between bacteria and phage OGUs, presence of experimentally validated protein family domains (Pfams) between phage and bacteria OGUs, and CRISPR targeting of bacteria toward phages (CRISPR). (C) Proportions of samples included (gray) and excluded (red) in the model. Samples were excluded from the model because they did not yield any scores. Those interactions without scores were automatically classified as not having interactions. (D) Network diameter (measure of graph size; the greatest number of traversed vertices required between two vertices), (E) number of vertices, and (F) number of edges (relationships) for the total network (orange) and the individual study sub-networks (diet study = red, skin study = yellow, twin study = green).

We used our random forest model to classify the relationships between bacteria and phage operational genomic units, which were then used to build the interactive network. The master network, analogous to the universal microbiome network concept previously described [57], contained the three studies as sub-networks, which themselves each contained sub-networks for each sample (**Supplemental Figure S9**).

Metadata including study, sample ID, disease, and OGU abundance within the community were stored in the master network for parsing in downstream analyses (**Supplemental Figure S9**). The phage and bacteria of the master network demonstrated both narrow broad ranges of infectious interactions (**Supplemental Figure S10**). Bacterial and phage relative abundance was recorded in each sample for each OGU and the weight of the edge connecting those OGUs was calculated as a function of those relative abundance values. The separate extraction of the phage and bacterial libraries ensured a more accurate measurement of the microbial communities, as has been outlined previously [58,59]. The master network was highly connected and contained 38,337 infectious relationships among 435 nodes, representing 155 phages and 280 bacteria. Although the network was highly connected, not all relationships were present in all samples. Relationships were weighted by the relative abundances of their associated bacteria and phages. Like the master network, the skin network exhibited a diameter of 4 (measure of graph size; the greatest number of traversed vertices required between two vertices) and included 433 (154 phages, 279 bacteria, 99.5% total) and 38,099 (99.4%) of the master network nodes and edges, respectively (**Figure 1 E-F**). Additionally, the subnetworks demonstrated narrow ranges of eccentricity across their nodes (**Supplemental Figure S11**). The phages and bacteria in the diet and twin sample sets were more sparsely related, with the diet study consisting of 80 (32 phages, 48 bacteria) nodes and 1,290 relationships, and the twin study containing 130 (29 phages, 101 bacteria) nodes and 2,457 relationships (**Figure 1 E-F**). As a validation measure, we identified five (1.7%) examples of phage OGUs which contained similar genomic elements to the four previously described, broadly infectious phages isolated from Lake Michigan (tblastx; e-value < 10^−25^) [60].

### Role of Diet on Gut Microbiome Connectivity

Diet is a major environmental factor that influences resource availability and gut microbiome composition and diversity, including bacteria and phages [14,61,62]. Previous work in isolated culture-based systems has suggested that changes in nutrient availability are associated with altered phage-bacteria network structures [30], although this has yet to be tested in humans. We therefore hypothesized that a change in diet would also be associated with a change in virome-microbiome network structure in the human gut.

We evaluated the diet-associated differences in gut virome-microbiome network structure by quantifying how central each sample’s network was on average. We accomplished this by utilizing two common weighted centrality metrics: degree centrality and closeness centrality. Degree centrality, the simplest centrality metric, was defined as the number of connections each phage made with each bacterium. We supplemented measurements of degree centrality with measurements of closeness centrality. Closeness centrality is a metric of how close each phage or bacterium is to all of the other phages and bacteria in the network. A higher closeness centrality suggests that the effects of genetic information or altered abundance would be more impactful to all other microbes in the system. Because these are weighted metrics, the values are functions of both connectivity as well as community composition. A network with higher average closeness centrality also indicates an overall greater degree of connections, which suggests a greater resilience against network degradation by extinction events [30,63]. This is because more highly connected networks are less likely to degrade into multiple smaller networks when bacteria or phages are randomly removed [30,63]. We used this information to calculate the average connectedness per sample, which was corrected for the maximum potential degree of connectedness. Unfortunately our dataset was insufficiently powered to make strong conclusions toward this hypothesis, but this is an interesting observation that warrants further investigation. This observation also serves to illustrate the types of questions we can answer with more comprehensive microbiome sampling and integrative analyses.

Using our small sample set, we observed that the gut microbiome network structures associated with high-fat diets appeared less connected than those of low-fat diets, although a greater sample size will be required to more properly evaluate this trend (**Figure 2 A-B**). Five subjects were available for use, all of which had matching bacteria and virome datasets and samples from 8-10 days following the initiation of their diets. High-fat diets appeared to exhibit reduced degree centrality (**Figure 2 A**), suggesting bacteria in high-fat environments were targeted by fewer phages and that phage tropism was more restricted. High-fat diets also appeared to exhibit decreased closeness centrality (**Figure 2 B**), indicating that bacteria and phages were more distant from other bacteria and phages in the community. This would make genetic transfer and altered abundance of a given phage or bacterium less capable of impacting other bacteria and phages within the network.

**Figure 2:**
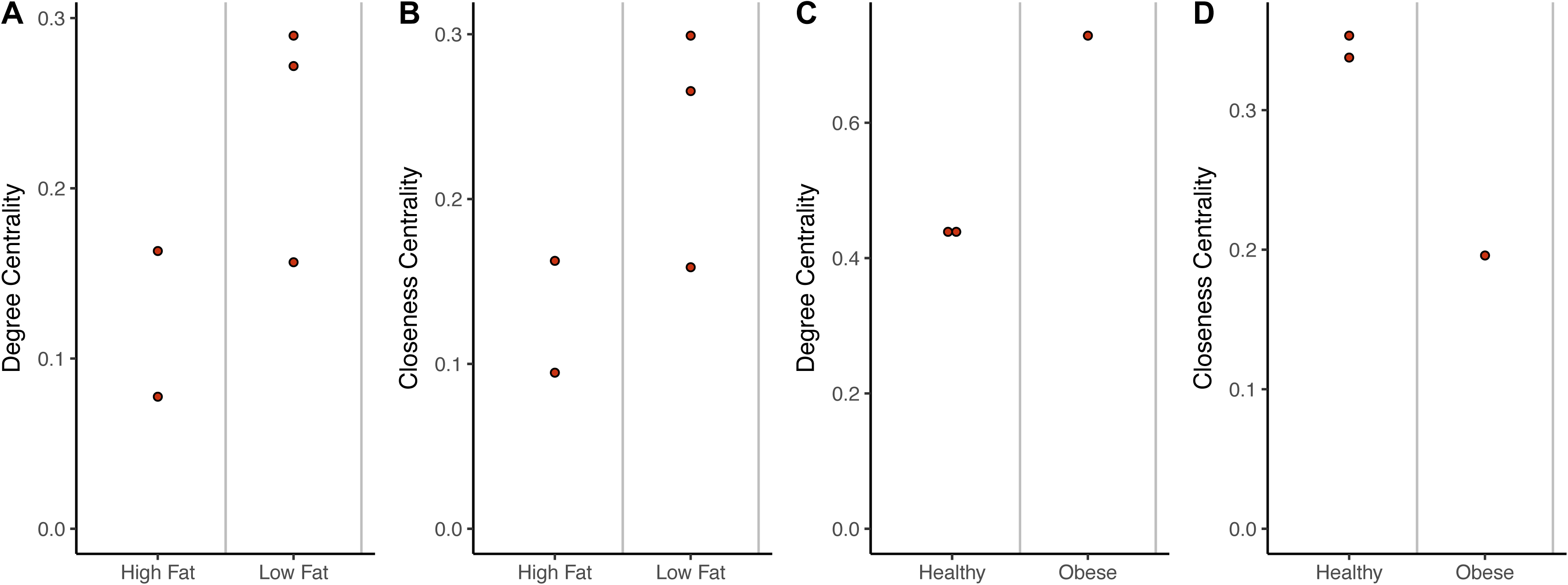
Impact of Diet and Obesity on Gut Network Structure. (A) Quantification of average degree centrality (number of edges per node) and (B) closeness centrality (average distance from each node to every other node) of gut microbiome networks of subjects limited to exclusively high-fat or low-fat diets. Each point represents the centrality from a human subject stool sample that was collected 8-10 days following the beginning of their defined diet. There are five samples here, compared to the four in figure 3, because one of the was only sampled post-diet, providing us data for this analysis but not allowing us to compare to a baseline for figure 3. (C) Quantification of average degree centrality and (D) closeness centrality between obese and healthy adult women from the Twin gut study. Each point represents a stool sample taken from one of the three adult woman confirmed as obese or healthy and with matching virus and bacteria data.

In addition to diet, we observed a possible trend that obesity influenced network structure. This was done using the three mother samples available from the twin sample set, all of which had matching bacteria and phage samples and confirmed BMI information. The obesity-associated network appeared to have a higher degree centrality (**Figure 2 C**), but less closeness centrality than the healthy-associated networks (**Figure 2 D**). These results suggested that the obesity-associated networks may be less connected. This again comes with the caveat that this is only an opportunistic observation using an existing sample set with too few samples to make more substantial claims. We included this observation as a point of interest, given the data was available.

### Individuality of Microbial Networks

Skin and gut community membership and diversity are highly personal, with people remaining more similar to themselves than to other people over time [13,64,65]. We therefore hypothesized that this personal conservation extended to microbiome network structure. We addressed this hypothesis by calculating the degree of dissimilarity between each subject’s network, based on phage and bacteria abundance and centrality. We quantified phage and bacteria centrality within each sample graph using the weighted eigenvector centrality metric. This metric defines central phages as those that are highly abundant (*A*_*O*_ as defined in the methods) and infect many distinct bacteria which themselves are abundant and infected by many other phages. Similarly, bacterial centrality was defined as those bacteria that were both abundant and connected to numerous phages that were themselves connected to many bacteria. We then calculated the similarity of community networks using the weighted eigenvector centrality of all nodes between all samples. Samples with similar network structures were interpreted as having similar capacities for network robustness and transmitting genetic material.

We used this network dissimilarity metric to test whether microbiome network structures were more similar within people than between people over time. We found that gut microbiome network structures clustered by person (ANOSIM p-value = 0.008, R = 1, **Figure 3 A**). Network dissimilarity within each person over the 8-10 day sampling period was less than the average dissimilarity between that person and others, although this difference was not statistically significant (p-value = 0.125, **Figure 3 B**). Four of the five available subjects were used because one of the subjects was not sampled at the initial time point. The lack of statistical confidence was likely due to the small sample size of this dataset.

**Figure 3:**
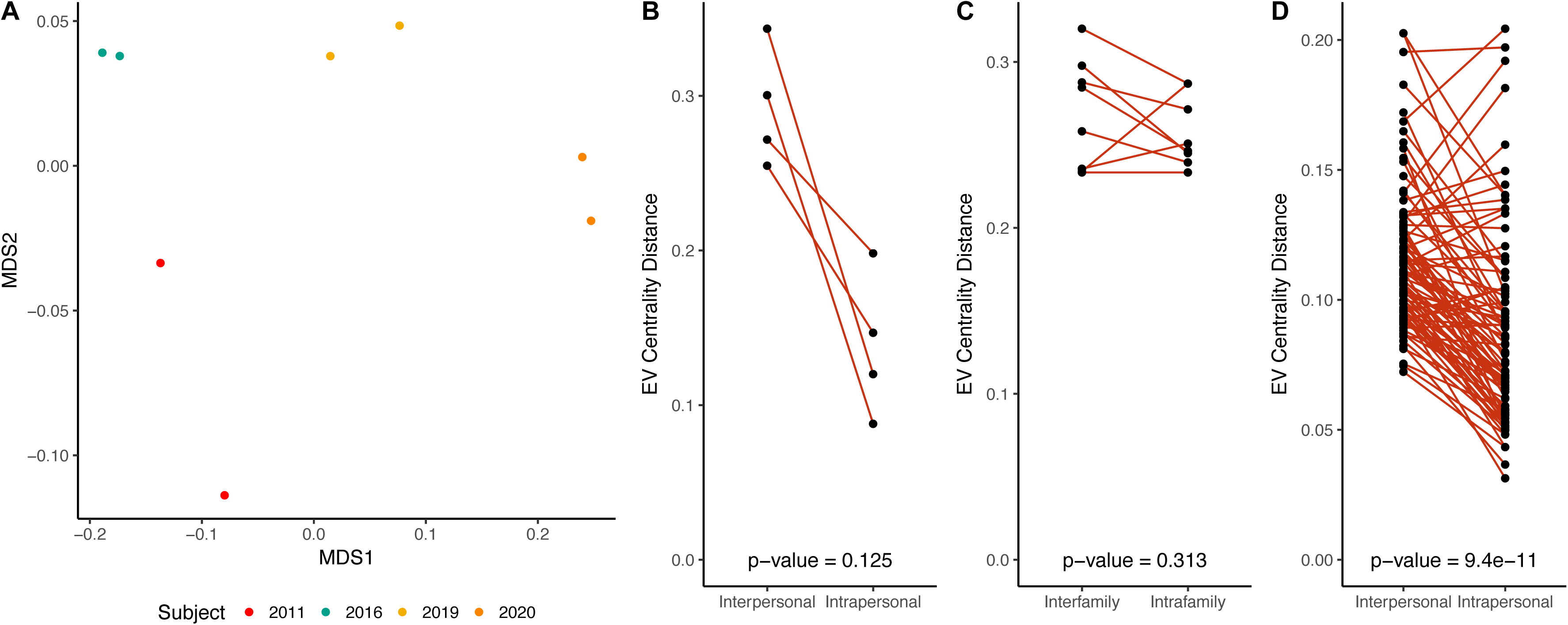
Intrapersonal vs Interpersonal Network Dissimilarity Across Different Human Systems. (A) NMDS ordination illustrating network dissimilarity between subjects over time. Each sample is colored by subject, with each colored sample pair collected 8-10 days apart. Dissimilarity was calculated using the Bray-Curtis metric based on abundance weighted eigenvector centrality signatures, with a greater distance representing greater dissimilarity in bacteria and phage centrality and abundance. Only four subjects were included here, compared to the five used in figure 2, because one of the subjects was missing the initial sampling time point and therefore lacked temporal sampling. (B) Quantification of gut network dissimilarity within the same subject over time (intrapersonal) and the mean dissimilarity between the subject of interest and all other subjects (interpersonal). The p-value is provided near the bottom of the figure. (C) Quantification of gut network dissimilarity within subjects from the same family (intrafamily) and the mean dissimilarity between subjects within a family and those of other families (interfamily). Each point represents the inter-family and intra-family dissimilarity of a twin or mother that was sampled over time. (D) Quantification of skin network dissimilarity within the same subject and anatomical location over time (intrapersonal) and the mean dissimilarity between the subject of interest and all other subjects at the same time and the same anatomical location (interpersonal). All p-values were calculated using a paired Wilcoxon test.

Although there was evidence for gut network conservation among individuals, we found no evidence for conservation of gut network structures within families. The gut network structures were not more similar within families (twins and their mothers; intrafamily) compared to other families (other twins and mothers; inter-family) (p-value = 0.547, **Figure 3 C**). In addition to the gut, skin microbiome network structure was conserved within individuals (p-value < 0.001, **Figure 3 D**). This distribution was similar when separated by anatomical sites. Most sites were statistically significantly more conserved within individuals (**Supplemental Figure S12**).

As an additional validation measure, we evaluated the tolerance of these findings to inaccuracies in the underlying network. As described above, our model is not perfect and there is likely to be noise from false positive and false negative predictions. We found that additional random noise, both by creating a fully connected graph or randomly reducing the number of edges to 60% of the original, changed the statistical significance values (p-values) of our findings but not by enough to change whether they were statistically significant (p-value < 0.05). Therefore the comparisons between groups are resilient to potential noise resulting from model false positive and false negative predictions (**Supplemental Figure S13**).

### Network Structures Across the Human Skin Landscape

Extensive work has illustrated differences in diversity and composition of the healthy human skin microbiome between anatomical sites, including bacteria, virus, and fungal communities [13,47,64]. These communities vary by degree of skin moisture, oil, and environmental exposure; features which were defined in the original publication [13]. As viruses are known to influence microbial diversity and community composition, we hypothesized that these differences would still be evident after integrating the bacterial and viral datasets and evaluating their microbe-virus network structure between anatomical sites. To test this, we evaluated the changes in network structure between anatomical sites within the skin dataset. The anatomical sites and their features (e.g. moisture & occlusion) were defined in the previous publication through consultation with dermatologists and reference to previous literature [13].

The average centrality of each sample was quantified using the weighted eigenvector centrality metric. Intermittently moist skin sites (dynamic sites that fluctuate between being moist and dry) were significantly more connected than the moist and sebaceous environments (p-value < 0.001, **Figure 4 A**). Also, skin sites that were occluded from the environment were less connected than those that were constantly exposed to the environment or only intermittently occluded (p-value < 0.001, **Figure 4 B**). We also confirmed that addition of noise to the underlying network, as described above, altered the values of statistical significance (p-values) but not by enough to change whether they were statistically significant (**Supplemental Figure S14**).

**Figure 4:**
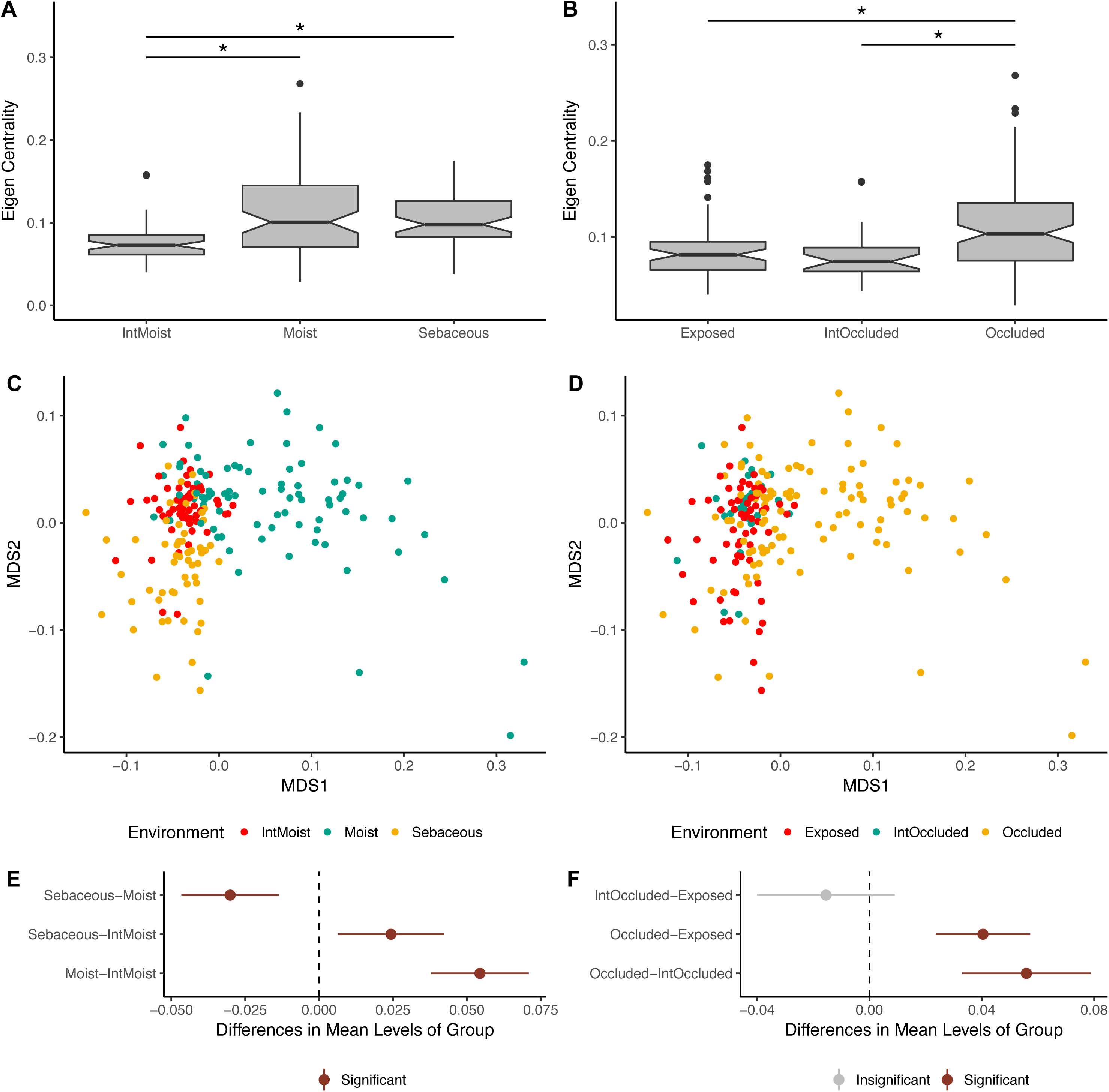
Impact of Skin Micro-Environment on Microbiome Network Structure. (A) Notched box-plot depicting differences in average eigenvector centrality between moist, intermittently moist, and sebaceous skin sites and (B) occluded, intermittently occluded, and exposed sites. Notched box-plots were created using ggplot2 and show the median (center line), the inter-quartile range (IQR; upper and lower boxes), the highest and lowest value within 1.5* IQR (whiskers), outliers (dots), and the notch which provides an approximate 95% confidence interval as defined by 1.58* IQR / sqrt(n). Sample sizes for each group were: Moist = 81, Sebaceous = 56, IntMoist = 56, Occluded = 106, Exposed = 61, IntOccluded = 26. (C) NMDS ordination depicting the differences in skin microbiome network structure between skin moisture levels and (D) occlusion. Samples are colored by their environment and their dissimilarity to other samples was calculated as described in figure 3. (E) The statistical differences of networks between moisture and (F) occlusion status were quantified with an anova and post hoc Tukey test. Cluster centroids are represented by dots and the extended lines represent the associated 95% confidence intervals. Significant comparisons (p-value < 0.05) are colored in red, and non-significant comparisons are gray.

To supplement this analysis, we compared the network signatures using the centrality dissimilarity approach described above. The dissimilarity between samples was a function of shared relationships, degree of centrality, and bacteria/phage abundance. When using this supplementary approach, we found that network structures significantly clustered by moisture, sebaceous, and intermittently moist status (**Figure 4 C,E**). Occluded sites were significantly different from exposed and intermittently occluded sites, but there was no difference between exposed and intermittently occluded sites (**Figure 4 D,F**). These findings provide further support that skin microbiome network structure differs significantly between skin sites.

## Discussion

Foundational work has provided a baseline understanding of the human microbiome by characterizing bacterial and viral diversity across the human body [13,14,46–48,66]. Here we integrated the bacterial and viral sequence sets to offer an initial understanding of how phage-bacteria networks differ throughout the human body, so as to provide a baseline for future studies of how and why microbiome networks differ in disease states. We implemented a network-based analytical model to evaluate the basic properties of the human microbiome through bacteria and phage relationships, instead of membership or diversity alone.

This approach enabled the application of network theory to provide a new perspective while analyzing bacterial and viral communities simultaneously. We utilized metrics of connectivity to model the extent to which communities of bacteria and phages interact through mechanisms such as horizontal gene transfer, modulated bacterial gene expression, and alterations in abundance.

Just as gut microbiome and virome composition and diversity are conserved in individuals [13,46,47,65], gut and skin microbiome network structures were conserved within individuals over time. Gut network structure was not conserved among family members. These findings suggested that the community properties inferred from microbiome interaction network structures, such as robustness (meaning a more highly connected network is more “robust” to network degradation because a randomly removed bacteria or phage node is less likely to divide or disintegrate [30,63] the overall network), the potential for horizontal gene transfer between members, and co-evolution of populations, were person-specific. These properties may be impacted by personal factors ranging from the body’s immune system to external environmental conditions, such as climate and diet.

We observed evidence supporting the ability of environmental conditions to shape gut and skin microbiome interaction network structure by observing that diet and skin location were associated with altered network structures. We observed evidence that diet was sufficient to alter gut microbiome network connectivity, although this needs to be interpreted cautiously as a case observation, due to the small sample size. Although the available sample size was small, our findings provide some preliminary evidence that high-fat diets are less connected than low-fat diets and that high-fat diets may therefore lead to less robust communities with a decreased ability for microbes to directly influence one another. We supported this finding with the observation that obesity may have been associated with decreased network connectivity. Together these findings suggest the food we eat may not only impact which microbes colonize our guts, but may also impact their interactions with infecting phages. Further work will be required to characterize these relationships with a larger cohort.

In addition to diet, the skin environment also influenced the microbiome interaction network structure. Network structure differed between environmentally exposed and occluded skin sites. The sites under greater environmental fluctuation and exposure (the exposed and intermittently exposed sites) were more connected and therefore were predicted to have a higher resilience against network degradation when random nodes are removed from the network. Likewise, intermittently moist sites demonstrated higher connectedness than the moist and sebaceous sites. These findings agree with previous work that has shown that bacterial community networks differ by skin environment types [57]. Together these data suggested that body sites under greater degrees of fluctuation harbored more highly connected microbiomes that are potentially more robust to network disruption by extinction events. This points to a link between microbiome and environmental robustness toward network homeostasis and warrants further investigation.

While these findings take us an important step closer to understanding the microbiome through interspecies relationships, there are caveats and considerations to our findings. First, as with most classification models, the infection classification model developed and applied is only as good as its training set – in this case, the collection of experimentally-verified positive and negative infection data. Large-scale experimental screens for phage and bacteria infectious interactions that report high-confidence negative interactions (i.e., no infection) are desperately needed, as they would provide more robust model training and improved model performance. Furthermore, just as we have improved on previous modeling efforts, we expect that new and creative scoring metrics will improve future performance. Other creative and high performing models are currently being developed and the applications of these models to community network creation will continue to move this field forward [43–45].

Second, although our analyses utilized the best datasets currently available for our study, this work was done retrospectively and relied on existing data up to seven years old. These archived datasets were limited by the technology and costs of the time. For example, the diet and twin studies, relied on multiple displacement amplification (MDA) in their library preparations–an approach used to overcome the large nucleic acids requirements typical of older sequencing library generation protocols. It is now known that MDA results in biases in microbial community composition [67], as well as toward ssDNA viral genomes [68,69], thus rendering the resulting microbial and viral metagenomes largely non-quantitative. Future work that employs larger sequence datasets and that avoids the use of bias-inducing amplification steps will build on and validate our findings, as well as inform the design and interpretation of further studies.

Although our models demonstrated satisfactory accuracy and overall performance, it was important to interpret our findings under the realization that our model was not perfect. This caveat is not new to the microbiome field, with a notable example being the use of 16S rRNA sequencing using the V4 variable region [59]. Use of the V4 variable region excluded detection of major skin bacterial members, meaning that the findings were not able to completely describe the underlying biological environment. Despite this caveat, skin microbiome studies provided valuable biological insights by focusing on the community differences between groups (e.g. disease and healthy) which were both analyzed the same way. Similarly, here we focused our conclusions on the differences between the groups which were all treated the same, so that we can minimize our dependence on a perfect predictive model. We also provided explicit evidence that the introduction of noise equally to the compared groups did not significantly impact our findings.

Third, the networks in this study were built using operational genomic units (OGUs), which represented groups of highly similar bacteria or phage genomes or clustered genome fragments. Similar clustering definition and validation methods, both computational and experimental, have been implemented in other metagenomic sequencing studies, as well [42,70–72]. These approaches could offer yet another level of sophistication to our network-based analyses. While this operationally defined clustering approach allows us to study whole community networks, our ability to make conclusions about interactions among specific phage or bacterial species or populations is inherently limited, compared to more focused, culture-based studies such as the work by Malki *et al* [60]. Future work must address this limitation, e.g., through improved binning methods and deeper metagenomic shotgun sequencing, but most importantly through an improved conceptual framing of what defines ecologically and evolutionarily cohesive units for both phage and bacteria [73]. Defining operational genomic units and their taxonomic underpinnings (e.g., whether OGU clusters represent genera or species) is an active area of work critical to the utility of this approach. As a first step, phylogenomic analyses have been performed to cluster cyanophage isolate genomes into informative groups using shared gene content, average nucleotide identity of shared genes, and pairwise differences between genomes [74]. Such population-genetic assessment of phage evolution, coupled with the ecological implications of genome heterogeneity, will inform how to define nodes in future iterations of the ecological network developed here. Even though we are hesitant to speculate on phage host ranges at low taxonomic levels in our dataset, the data does agree with previous reports of instances of broad phage host range [60,75]. Additionally, visualization of our dataset interactions using the heat map approach previously used in other host range studies, suggests a trend toward modular and nested tropism, but we do not have the strain-level resolution that powered those previous experimental studies.

Finally, it is important to note that our model was built using available full genomes with known interactions, while the experimental datasets resulted in OGUs created from metagenomic shotgun sequence sets, as described above. While this is an informative approach given available data, it is not ideal. We envision future work focusing on training models using metagenomic shotgun sample sets from “mock communities” of bacteria and phages with experimentally validated infectious relationships. This would also be more informative than relying on simulated metagenomic sample sets, whose use would result in models built on simulations and more assumptions instead of empirical data. Together this way the training set can be subjected to the same pre-processing, contig assembly, and OGU binning processes as the experimental data. Furthermore, exciting advances in long read sequencing platforms such as the Oxford Nanopore MinIon system will provide more accurate genomic scaffolds than *de novo* assembled contigs, allowing for more accurate training and predictions of our models. As discussed above, it is because our current model is susceptible to this noise that we focus our conclusions on comparisons between experimental groups that were both treated the same. This is also why it was important for us to evaluate the susceptibility of our results to noise caused by the less-than-perfect prediction model.

Together our work takes an initial step towards defining bacteria-virus interaction profiles as a characteristic of human-associated microbial communities. This approach revealed the impacts that different human environments (e.g., the skin and gut) can have on microbiome connectivity. By focusing on relationships between bacterial and viral communities, they are studied as the interacting cohorts they are, rather than as independent entities. While our developed bacteria-phage interaction framework is a novel conceptual advance, the microbiome also consists of archaea and small eukaryotes, including fungi and *Demodex* mites [1,76] – all of which can interact with human immune cells and other non-microbial community members [77]. Future work will build from our approach and include these additional community members and their diverse interactions and relationships (e.g., beyond phage-bacteria). This will result in a more robust network and a more holistic understanding of the evolutionary and ecological processes that drive the assembly and function of the human-associated microbiome.

## Materials & Methods

### Code Availability

A reproducible version of this manuscript written in R markdown and all of the code used to obtain and process the sequencing data is available at the following GitHub repository:

https://github.com/SchlossLab/Hannigan_ConjunctisViribus_ploscompbio_2017

### Data Acquisition & Quality Control

Raw sequencing data and associated metadata were acquired from the NCBI sequence read archive (SRA). Supplementary metadata were acquired from the same SRA repositories and their associated manuscripts. The gut virome diet study (`SRA: SRP002424`), twin virome studies (`SRA: SRP002523; SRP000319`), and skin virome study (SRA: SRP049645) were downloaded as. `sra` files. For clarity, the sample sizes used for each study subset were described with the data in the results section. Sequencing files were converted to `fastq` format using the `fastq-dump` tool of the NCBI SRA Toolkit (v2.2.0). Sequences were quality trimmed using the Fastx toolkit (v0.0.14) to exclude bases with quality scores below 33 and shorter than 75 bp [78]. Paired end reads were filtered to exclude sequences missing their corresponding pair using the `get_trimmed_pairs.py` script available in the source code.

### Contig Assembly

Contigs were assembled using the Megahit assembly program (v1.0.6) [79]. A minimum contig length of 1 kb was used. Iterative k-mer stepping began at a minimum length of 21 and progressed by 20 until 101. All other default parameters were used.

Contig simulations were performed by randomly extracting a string of genomic nucleotides that represented a defined percent length of that genome. This was accomplished using our RandomContigGenerator.pl, which was published in the associated GitHub repository.

### Contig Abundance Calculations

Contigs were concatenated into two master files prior to alignment, one for bacterial contigs and one for phage contigs. Sample sequences were aligned to phage or bacterial contigs using the Bowtie2 global aligner (v2.2.1) [80]. We defined a mismatch threshold of 1 bp and seed length of 25 bp. Sequence abundance was calculated from the Bowtie2 output using the `calculate_abundance_from_sam.pl` script available in the source code.

### Operational Genomic Unit Binning

Contigs often represent large fragments of genomes. In order to reduce redundancy and the resulting artificially inflated genomic richness within our dataset, it was important to bin contigs into operational units based on their similarity. This approach is conceptually similar to the clustering of related 16S rRNA sequences into operational taxonomic units (OTUs), although here we are clustering contigs into operational genomic units (OGUs) [66].

Contigs were clustered using the CONCOCT algorithm (v0.4.0) [81]. Because of our large dataset and limits in computational efficiency, we randomly subsampled the dataset to include 25% of all samples, and used these to inform contig abundance within the CONCOCT algorithm. CONCOCT was used with a maximum of 500 clusters, a k-mer length of four, a length threshold of 1 kb, 25 iterations, and exclusion of the total coverage variable.

OGU abundance (*A*_*O*_) was obtained as the sum of the abundance of each contig (*A*_*j*_) associated with that OGU. The abundance values were length corrected such that:

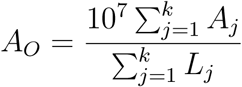

Where L is the length of each contig j within the OGU.

### Operational Genomic Unit Identification

To confirm a lack of phage sequences in the bacterial OGU dataset, we performed blast nucleotide alignment of the bacterial OGU representative sequences using an e-value < 10^−25^, which was stricter than the 10^−10^ threshold used in the random forest model below, against all of the phage reference genomes available in the EMBL database. We used a stricter threshold because we know there are genomic similarities between bacteria and phage OGUs from the interactive model, but we were interested in contigs with high enough similarity to references that they may indeed be from phages. We also performed the converse analysis of aligning phage OGU representative sequences to EMBL bacterial reference genomes. We ran both the phage and bacteria OGU representative sequences through the Virsorter program (1.0.3) to identify phages (all default parameters were used), using only those in the high confidence identification category “class 1” [50]. Finally, we filtered out phage OGUs that had bacterial elements as described above, but also lacked known phage elements by using the tblastx algorithm and a maximum e-value of 10^−25^.

### Open Reading Frame Prediction

Open reading frames (ORFs) were identified using the Prodigal program (V2.6.2) with the meta mode parameter and default settings [82].

### Classification Model Creation and Validation

The classification model for predicting interactions was built using experimentally validated bacteria-phage infections or validated lack of infections from six studies [41,51–55]. No further reference databases were used in our alignment procedures. Associated reference genomes were downloaded from the European Bioinformatics Institute (see details in source code). The model was created based on the four metrics listed below.

The four scores were used as parameters in a random forest model to classify bacteria and bacteriophage pairs as either having infectious interactions or not. The classification model was built using the Caret R package (v6.0.73) [83]. The model was trained using five-fold cross validation with ten repeats, and the median model performance was evaluated by training the model on 80% of the dataset and testing performance on the remaining 20%. Pairs without scores were classified as not interacting. The model was optimized using the ROC value. The resulting model performance was plotted using the plotROC R package.

### Identify Bacterial CRISPRs Targeting Phages

Clustered Regularly Interspaced Short Palindromic Repeats (CRISPRs) were identified from bacterial genomes using the PilerCR program (v1.06) [84]. Resulting spacer sequences were filtered to exclude spacers shorter than 20 bp and longer than 65 bp. Spacer sequences were aligned to the phage genomes using the nucleotide BLAST algorithm with default parameters (v2.4.0) [85]. The mean percent identity for each matching pair was recorded for use in our classification model.

### Detect Matching Prophages within Bacterial Genomes

Temperate bacteriophages infect and integrate into their bacterial host’s genome. We detected integrated phage elements within bacterial genomes by aligning phage genomes to bacterial genomes using the nucleotide BLAST algorithm and a minimum e-value of 1e-10. The resulting bitscore of each alignment was recorded for use in our classification model.

### Identify Shared Genes Between Bacteria and Phages

As a result of gene transfer or phage genome integration during infection, phages may share genes with their bacterial hosts, providing us with evidence of phage-host pairing. We identified shared genes between bacterial and phage genomes by assessing amino acid similarity between the genes using the Diamond protein alignment algorithm (v0.7.11.60) [86]. The mean alignment bitscores for each genome pair were recorded for use in our classification model.

### Protein-Protein Interactions

The final method used for predicting infectious interactions between bacteria and phages was the detection of pairs of genes whose proteins are known to interact. We assigned bacterial and phage genes to protein families by aligning them to the Pfam database using the Diamond protein alignment algorithm. We then identified which pairs of proteins were predicted to interact using the Pfam interaction information within the Intact database [56]. The mean bitscores of the matches between each pair were recorded for use in the classification model.

### Secondary Dataset Validation

The performance of our model for identifying diverse infectious relationships between bacteria and phages, beyond those that were included in the model creation step, were validated using additional bacterial and phage reference genomes, which could be linked by the records of which phage strains were isolated on which bacteria under laboratory conditions. Viral and bacterial reference genomes were downloaded from the GenBank repository on February 19, 2018 using the viral location ftp://ftp.ncbi.nih.gov/refseq/release/vira and the bacterial location ftp://ftp.ncbi.nih.gov/refseq/release/bacteria/. This resulted in the use of 539 complete phages reference genomes (with identified hosts) and 3,469 bacterial reference genomes. We used the same prediction model to predict which phages were infecting which hosts, so as to confirm that the model was capable of identifying interactions in a more diverse dataset. Bacteria interactions were identified at the species level. The random contig iteration analysis was performed using a subset of bacterial reference genomes, for computational performance reasons. Only single representative genomes for each species were used.

### Interaction Network Construction

The bacteria and phage operational genomic units (OGUs) were scored using the same approach as outlined above. The infectious pairings between bacteria and phage OGUs were classified using the random forest model described above. The predicted infectious pairings and all associated metadata were used to populate a graph database using Neo4j graph database software (v2.3.1) [87]. This network was used for downstream community analysis. Tolerance to false negative and false positive noise within the networks was assessed by randomly removing a defined fraction of network edges before re-running the downstream analysis work flows. This was accomplished using functionality within the igraph R package (v1.0.1) [88].

### Centrality Analysis

We quantified the centrality of graph vertices using three different metrics, each of which provided different information graph structure. When calculating these values, let *G*(*V, E*) be an undirected, unweighted graph with *|V |* = *n* nodes and *|E|* = *m* edges. Also, let **A** be its corresponding adjacency matrix with entries *a*_*ij*_ = 1 if nodes *V*_*i*_ and *V*_*j*_ are connected via an edge, and *a*_*ij*_ = 0 otherwise.

Briefly, the **closeness centrality** of node *V*_*i*_ is calculated taking the inverse of the average length of the shortest paths (d) between nodes *V*_*i*_ and all the other nodes *V*_*j*_. Mathematically, the closeness centrality of node *V*_*i*_ is given as:

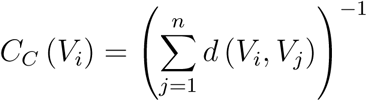

The distance between nodes (d) was calculated as the shortest number of edges required to be traversed to move from one node to another.

Intuitively, the **degree centrality** of node *V*_*i*_ is defined as the number of edges that are incident to that node:

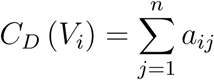

where *a*_*ij*_ is the *ij*^*th*^ entry in the adjacency matrix **A**.

The eigenvector centrality of node *V*_*i*_ is defined as the *i*^*th*^ value in the first eigenvector of the associated adjacency matrix **A**. Conceptually, this function results in a centrality value that reflects the connections of the vertex, as well as the centrality of its neighboring vertices.

The **centralization** metric was used to assess the average centrality of each sample graph **G**. Centralization was calculated by taking the sum of each vertex *V*_*i*_’s centrality from the graph maximum centrality *C*_*w*_, such that:

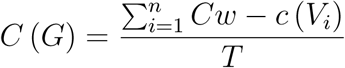

The values were corrected for uneven graph sizes by dividing the centralization score by the maximum theoretical centralization (T) for a graph with the same number of vertices.

Degree and closeness centrality were calculated using the associated functions within the igraph R package (v1.0.1) [88].

### Network Relationship Dissimilarity

We assessed similarity between graphs by evaluating the shared centrality of their vertices, as has been done previously. More specifically, we calculated the dissimilarity between graphs *G*_*i*_ and *G*_*j*_ using the Bray-Curtis dissimilarity metric and eigenvector centrality values such that:

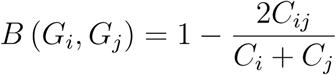

Where *C*_*ij*_ is the sum of the lesser centrality values for those vertices shared between graphs, and *C*_*i*_ and *C*_*j*_ are the total number of vertices found in each graph. This allows us to calculate the dissimilarity between graphs based on the shared centrality values between the two graphs.

### Statistics and Comparisons

Differences in intrapersonal and interpersonal network structure diversity, based on multivariate data, were calculated using an analysis of similarity (ANOSIM). Statistical significance of univariate Eigenvector centrality differences were calculated using a paired Wilcoxon test.

Statistical significance of differences in univariate eigenvector centrality measurements of skin virome-microbiome networks were calculated using a pairwise Wilcoxon test, corrected for multiple hypothesis tests using the Holm correction method. Multivariate eigenvector centrality was measured as the mean differences between cluster centroids, with statistical significance measured using an ANOVA and post hoc Tukey test.

## Acknowledgments

We thank the members of the Schloss lab for their underlying contributions. We thank the authors of the original studies for making their data and metadata publicly available and understandable. We also thank the participants in the studies.

## Author Contributions

*Conceptualization*: GDH, MBD, DK, PDS. *Data Curation*: GDH. *Formal Analysis*: GDH. *Funding Acquisition*: GDH, PDS. *Writing – Original Draft Preparation*: GDH, PDS. *Writing – Review & Editing*: GDH, MBD, DK, PDS.

## Supplemental Figure Captions

Figure S1: **Sequencing Depth Summary.** Number of sequences that aligned to (A) Phage and (B) Bacteria operational genomic units per sample and colored by study.

Figure S2: **Contig Summary Statistics.** Scatter plot heat map with each hexagon representing the abundance of contigs. Contigs are organized by length on the x-axis and the number of aligned sequences on the y-axis.

Figure S3: **Operational Genomic Unit Summary Statistics.** Scatter plot with operational genomic unit clusters organized by average contig length within the cluster on the x-axis and the number of contigs in the cluster on the y-axis. Operational genomic units of (A) bacteriophages and (B) bacteria are shown.

Figure S4: **Summary information of validation dataset used in the interaction predictive model.** A) Categorical heat-map highlighting the experimentally validated positive and negative interactions. Only bacteria species are shown, which represent multiple reference strains. Phages are labeled on the x-axis and bacteria are labeled on the y-axis. B) Quantification of bacterial host strains known to exist for each phage. C) Genome strandedness and D) linearity of the phage reference genomes used for the dataset.

Figure S5: **Ability of prediction model to identify broad range of bacteria and phage interactions.** Each complete bacteriophage labeled on the y axis. The number of complete bacterial species genomes that were correctly predicted to be infected by each phage, according to the GenBank records, is on the x-axis. True positive interactions are colored in purple, and false negative interactions are yellow. Because this dataset only had confirmed interactions and not confirmed lack of interactions, false positive and true negative values could be not determined.

Figure S6: **Impact of incomplete genomic sequences on model performance.** Number of correctly identified infectious interactions between bacteria and phages are presented on the y-axis. The fraction of genomic length that was used to randomly extract contigs from reference bacterial and phage sequences is presented on the x-axis (e.g. 0.9 means contig lengths were 90% of the total genome length). Overall this presents a quantification of the loss of identified infectious relationships as the percent of available genomic material is reduced.

Figure S7: **Classification Model Performance By Nested Cross-Validation.** Box plot illustrating the median and variance of phage-bacteria interaction prediction model. Performance was evaluated using nested cross validation, meaning that 20% of the samples were randomly withheld from model training and then used to evaluate performance. The results of 100 random iterations are shown. Metrics include area under the curve (gray), sensitivity (red), and specificity (tan).

Figure S8: **Stable Classification Model Performance Over Random Iterations** In addition to nested cross-validation, here we show the results from the five-fold cross validation, in which 20% of the samples were randomly withheld during the training stage for model evaluation and mtry tuning (parameter defined in the R Random Forest package, which is implemented in Caret, as “the number of variables randomly sampled as candidates at each split”). The results of 25 random iterations are shown. Metrics include area under the curve (red), sensitivity (green), and specificity (blue). Dashed line highlight the random point of 0.5.

Figure S9: **Structure of the interactive network.** Metadata relationships to samples (Phage Sample ID and Bacteria Sample ID) included the associated time point, the study, the subject the sample was taken from, and the associated disease. Infectious interactions were recorded between phage and bacteria operational genomic units (OGUs). Sequence count abundance for each OGU within each sample was also recorded.

Figure S10: **Heatmap of Phage-Bacteria Interaction Relationships of Master Network.** Heatmap illustrating the ranges of infectious interactions predicted between bacteria and bacteriophages across our three studies. Bacterial OGUs are aligned on the vertical access, and the bacteriophage OGUs are organized on the horizontal access. OGUs are organized near other OGUs with similar infectious profiles, which are further illustrated by the dendrograms. Predicted infections are tan and predicted lacks of interactions are red.

Figure S11: **Distribution of node eccentricity across subnetworks.** Histograms illustrating the distributions of node eccentricity values across the subnetworks, for supplementing the node, edge, and diameter values provided for the networks. Eccentricity of each node is the shortest distance of that node to the furthest other node within the graph.

Figure S12: **Intrapersonal vs Interpersonal Dissimilarity of the Skin.** Quantification of skin network dissimilarity within the same subject and anatomical location over time (intrapersonal) and the mean dissimilarity between the subject of interest and all other subjects at the same time and the same anatomical location (interpersonal), separated by each anatomical site (forehead [Fh], palm [Pa], toe web [Tw], umbilicus [Um], antecubital fossa [Ac], axilla [Ax], and retroauricular crease [Ra]). P-value was calculated using a paired Wilcoxon test.

Figure S13: **P-values of interpersonal group diversity differences with graph edge noise.** The x-axis represents the percent of errors that were randomly removed (or added) as a means to evaluate the impact of random noise on statistical significance of group differences. Resulting p-values for each graph is shown on the y-axis. The dot and bars are the mean and standard error of five iterations of group comparisons with random edge removal. Significance of A) ANOSIM p-value of diet network dissimilarity, B) p-value of interpersonal and intrapersonal diet network dissimilarity, C) p-value of interpersonal and intrapersonal skin network dissimilarity, and D) p-value of interpersonal and intrapersonal twin gut network dissimilarity. This corresponds to the findings in Figure 3.

Figure S14: **P-values of differences in Eigen Centrality between skin site microbiome networks.** The x-axis represents the percent of errors that were randomly removed (or added) as a means to evaluate the impact of random noise on statistical significance of group differences. Resulting p-values for each graph is shown on the y-axis. The dot and bars are the mean and standard error of five iterations of group comparisons with random edge removal. The groups compared were the degrees of site moisture (left) and occlusion (right). The findings correspond to pannels A and B in Figure 4.

## Supplemental Table Captions

Table S1: Summary of the primary quality control measures reported in the original publications of the viromes used in this current study.

Table S2: The positive and negative bacteria and bacteriophage interactions used to train the prediction model, as also illustrated in Figure S4. Citation sources are also included.

