## Supplementary Materials for "Biogeography & environmental conditions shape bacteriophage-bacteria networks across the human microbiome"

**Table S1**

| Study | Citation | Virome Quality Control Measures |
| --- | --- | --- |
| Diet & the Gut Virome | Minot, 2011 | • 16S rRNA gene qPCR revealed reduction in bacterial DNA of at least 10,000X.<br>• Alignment of shotgun sequences revealed 35X reduction in 16S rRNA gene alignments in virome compared to bacteria shotgun.<br>• Electron microscopy and nucleic acid stain techniques visually confirmed lack of bacteria in virome samples. |
| Skin Virome | Hannigan, 2015 | • Significant reduction in reads mapping to 16S rRNA gene sequence, compared to bacteria shotgun dataset.<br>• Significant reduction in reads mapping to human genome, compared to bacteria shotgun dataset.<br>• Average viral relative abundance of 0.4% in bacterial shotgun dataset. |
| Twin Gut Virome | Reyes, 2010 | • Confirmation that 2.5% of bacterial shotgun reads mapped to virome, and 76% of virome reads matched the shotgun 2.5%. |
