## Supplementary Materials for "Biogeography & environmental conditions shape bacteriophage-bacteria networks across the human microbiome"

*Table S2*

| Bacterial Host | Bacteriophage | Interaction | Citation |
|---|---|---|---|
| Achromobacter xylosoxidans | Pbunalikevirus phiFenriz | Negative | Malki, *et al.* 2015. |
| Achromobacter xylosoxidans | Pbunalikevirus phiHabibi | Negative | Malki, *et al.* 2015. |
| Achromobacter xylosoxidans | Pbunalikevirus phiMoody | Negative | Malki, *et al.* 2015. |
| Achromobacter xylosoxidans | Pbunalikevirus phiVader | Negative | Malki, *et al.* 2015. |
| Acinetobacter baumannii | Pseudomonas phage PA1phi | Negative | Kim *et al.* 2012. |
| Acinetobacter baumannii | Acinetobacter phage Petty | Positive | Edwards *et al.* 2015. |
| Arthrobacter sp. | Pbunalikevirus phiHabibi | Positive | Malki, *et al.* 2015. |
| Arthrobacter sp. | Pbunalikevirus phiVader | Positive | Malki, *et al.* 2015. |
| Arthrobacter sp. | Pbunalikevirus phiMoody | Positive | Malki, *et al.* 2015. |
| Arthrobacter sp. | Pbunalikevirus phiFenriz | Positive | Malki, *et al.* 2015. |
| Arthrobacter sulfonivorans | Pbunalikevirus phiHabibi | Positive | Malki, *et al.* 2015. |
| Arthrobacter sulfonivorans | Pbunalikevirus phiVader | Positive | Malki, *et al.* 2015. |
| Arthrobacter sulfonivorans | Pbunalikevirus phiMoody | Positive | Malki, *et al.* 2015. |
| Arthrobacter sulfonivorans | Pbunalikevirus phiFenriz | Positive | Malki, *et al.* 2015. |
| Bacillus pumilus | Enterobacteria phage phi92 | Negative | Schwarzer, *et al.* 2012. |
| Bacillus pumilus | Enterobacteria phage K1F | Negative | Schwarzer, *et al.* 2012. |
| Bacillus amyloliquefaciens | Enterobacteria phage phi92 | Negative | Schwarzer, *et al.* 2012. |
| Bacillus amyloliquefaciens | Enterobacteria phage K1F | Negative | Schwarzer, *et al.* 2012. |
| Bacillus subtilis | Enterobacteria phage phi92 | Negative | Schwarzer, *et al.* 2012. |
| Bacillus subtilis | Enterobacteria phage K1F | Negative | Schwarzer, *et al.* 2012. |

| Bacterial Host | Bacteriophage | Interaction | Citation |
|---|---|---|---|
| Burkholderia cenocepacia | Burkholderia phage KL1 | Positive | Hargreaves, *et al.* 2014. |
| Burkholderia cenocepacia | Burkholderia phage KS14 | Positive | Edwards *et al.* 2015. |
| Chryseobacterium sp. | Pbunalikevirus phiHabibi | Positive | Malki, *et al.* 2015. |
| Chryseobacterium sp. | Pbunalikevirus phiFenriz | Positive | Malki, *et al.* 2015. |
| Chryseobacterium sp. | Pbunalikevirus phiMoody | Positive | Malki, *et al.* 2015. |
| Chryseobacterium sp. | Pbunalikevirus phiVader | Positive | Malki, *et al.* 2015. |
| Clostridium perfringens | Clostridium phage phiCP7R | Positive | Edwards *et al.* 2015. |
| Clostridium perfringens | Clostridium phage phiZP2 | Positive | Edwards *et al.* 2015. |
| Corynebacterium glutamicum | Corynebacterium phage P1201 | Positive | Edwards *et al.* 2015. |
| Cronobacter sakazakii | Cronobacter phage CR5 | Positive | Edwards *et al.* 2015. |
| Escherichia coli | Pbunalikevirus phiHabibi | Positive | Malki, *et al.* 2015. |
| Escherichia coli | Pbunalikevirus phiMoody | Positive | Malki, *et al.* 2015. |
| Escherichia coli | Pbunalikevirus phiFenriz | Positive | Malki, *et al.* 2015. |
| Escherichia coli | Pbunalikevirus phiVader | Positive | Malki, *et al.* 2015. |
| Escherichia coli | Pseudomonas aeruginosa phage F116 | Negative | Jensen *et al.* 1998. |
| Escherichia coli | Pseudomonas phage D3 | Negative | Jensen *et al.* 1998. |
| Enterobacter aerogenes | Pseudomonas phage PA1phi | Negative | Kim *et al.* 2012. |
| Erwinia amylovora | Erwinia phage phiEaH2 | Positive | Edwards *et al.* 2015. |
| Escherichia coli | Pseudomonas phage PA1phi | Negative | Kim *et al.* 2012. |
| Escherichia coli | Enterobacteria phage K1F | Positive | Schwarzer, *et al.* 2012. |
| Escherichia coli | Enterobacteria phage phi92 | Positive | Schwarzer, *et al.* 2012. |
| Klebsiella pneumoniae | Klebsiella phage KP15 | Positive | Edwards *et al.* 2015. |
| Lactococcus lactis | Lactococcus phage 949 | Positive | Edwards *et al.* 2015. |
| Lactococcus lactis | Lactococcus phage phiL47 | Positive | Edwards *et al.* 2015. |
| Listeria monocytogenes | Pseudomonas phage PA1phi | Positive | Kim *et al.* 2012. |
| Microbacterium sp. | Pbunalikevirus phiVader | Positive | Malki, *et al.* 2015. |
| Microbacterium sp. | Pbunalikevirus phiMoody | Positive | Malki, *et al.* 2015. |

| Bacterial Host | Bacteriophage | Interaction | Citation |
|---|---|---|---|
| Microbacterium sp. | Pbunalikevirus phiHabibi | Positive | Malki, *et al.* 2015. |
| Microbacterium sp. | Pbunalikevirus phiFenriz | Positive | Malki, *et al.* 2015. |
| Mycobacterium smegmatis | Mycobacterium phage SWU1 | Positive | Edwards *et al.* 2015. |
| Pseudomonas aeruginosa | Pbunalikevirus phiMoody | Positive | Malki, *et al.* 2015. |
| Pseudomonas aeruginosa | Pbunalikevirus phiVader | Positive | Malki, *et al.* 2015. |
| Pseudomonas aeruginosa | Pbunalikevirus phiFenriz | Positive | Malki, *et al.* 2015. |
| Pseudomonas aeruginosa | Pbunalikevirus phiHabibi | Positive | Malki, *et al.* 2015. |
| Pseudomonas aeruginosa | Pseudomonas aeruginosa phage F116 | Positive | Jensen *et al.* 1998. |
| Pseudomonas aeruginosa | Pseudomonas phage D3 | Positive | Jensen *et al.* 1998. |
| Photobacterium angustum | Vibrio phage KVP40 | Negative | Matsuza *et al.* 1992. |
| Photobacterium leiognathi | Vibrio phage KVP40 | Positive | Matsuza *et al.* 1992. |
| Propionibacterium acnes | Propionibacterium phage PHL112N00 | Positive | Edwards *et al.* 2015. |
| Pseudomonas aeruginosa | Pseudomonas phage PA1phi | Positive | Kim *et al.* 2012. |
| Pseudomonas putida | Enterobacteria phage K1F | Negative | Schwarzer, *et al.* 2012. |
| Pseudomonas putida | Enterobacteria phage phi92 | Negative | Schwarzer, *et al.* 2012. |
| Ralstonia solanacearum | Ralstonia phage RSM3 | Positive | Edwards *et al.* 2015. |
| Ralstonia solanacearum | Ralstonia phage RSS1 | Positive | Edwards *et al.* 2015. |
| Sphaerotilus natans | Pseudomonas aeruginosa phage F116 | Positive | Jensen *et al.* 1998. |
| Sphaerotilus natans | Pseudomonas phage D3 | Positive | Jensen *et al.* 1998. |
| Salmonella enterica | Pbunalikevirus phiFenriz | Negative | Malki, *et al.* 2015. |
| Salmonella enterica | Pbunalikevirus phiHabibi | Negative | Malki, *et al.* 2015. |
| Salmonella enterica | Salmonella phage FSL SP-058 | Positive | Edwards *et al.* 2015. |
| Salmonella enterica | Pbunalikevirus phiVader | Negative | Malki, *et al.* 2015. |
| Salmonella enterica | Pbunalikevirus phiMoody | Negative | Malki, *et al.* 2015. |
| Salmonella enterica | Enterobacteria phage phi92 | Positive | Schwarzer, *et al.* 2012. |

| Bacterial Host | Bacteriophage | Interaction | Citation |
|---|---|---|---|
| Salmonella enterica | Enterobacteria phage K1F | Negative | Schwarzer, *et al.* 2012. |
| Serratia marcescens | Pseudomonas phage PA1phi | Negative | Kim *et al.* 2012. |
| Shigella boydii | Pbunalikevirus phiVader | Negative | Malki, *et al.* 2015. |
| Shigella boydii | Pbunalikevirus phiHabibi | Negative | Malki, *et al.* 2015. |
| Shigella boydii | Pbunalikevirus phiFenriz | Negative | Malki, *et al.* 2015. |
| Shigella boydii | Pbunalikevirus phiMoody | Negative | Malki, *et al.* 2015. |
| Shigella flexneri | Pbunalikevirus phiFenriz | Negative | Malki, *et al.* 2015. |
| Shigella flexneri | Pbunalikevirus phiVader | Negative | Malki, *et al.* 2015. |
| Shigella flexneri | Pbunalikevirus phiMoody | Negative | Malki, *et al.* 2015. |
| Shigella flexneri | Pbunalikevirus phiHabibi | Negative | Malki, *et al.* 2015. |
| Shigella sonnei | Pseudomonas phage PA1phi | Positive | Kim *et al.* 2012. |
| Staphylococcus aureus | Pseudomonas phage PA1phi | Positive | Kim *et al.* 2012. |
| Staphylococcus epidermidis | Staphylococcus phage CNPH82 | Positive | Edwards *et al.* 2015. |
| Staphylococcus epidermidis | Pseudomonas phage PA1phi | Negative | Kim *et al.* 2012. |
| Streptococcus agalactiae | Pseudomonas phage PA1phi | Negative | Kim *et al.* 2012. |
| Streptococcus gordonii | Pseudomonas phage PA1phi | Negative | Kim *et al.* 2012. |
| Streptococcus pneumoniae | Streptococcus phage K13 | Positive | Edwards *et al.* 2015. |
| Streptococcus pneumoniae | Pseudomonas phage PA1phi | Negative | Kim *et al.* 2012. |
| Streptococcus salivarius | Pseudomonas phage PA1phi | Positive | Kim *et al.* 2012. |
| Thermus thermophilus | Thermus phage TMA | Positive | Edwards *et al.* 2015. |
| Vibrio natriegens | Vibrio phage KVP40 | Positive | Matsuza *et al.* 1992. |
| Vibrio parahaemolyticus | Vibrio phage KVP40 | Positive | Matsuza *et al.* 1992. |
| Vibrio vulnificus | Vibrio phage KVP40 | Negative | Matsuza *et al.* 1992. |
| Xylella fastidiosa | Xylella phage Xfas53 | Positive | Edwards *et al.* 2015. |
